# YAP restricts renal inflammation and mitigates kidney damage in nephronothisis related kidney disease

**DOI:** 10.1101/2022.01.17.475784

**Authors:** Giulia Ferri, Valentine Goffette, Esther Porée, Simone Braeg, Shana Carvalho, Marceau Quatredeniers, E. Wolfgang Kuehn, Sophie Saunier, Amandine Viau

## Abstract

Nephronophthisis (NPH) is an orphan recessive kidney disease mostly caused by mutations in *NPHP1* and 20 other genes encoding proteins that localize to primary cilia. To date the pathways linking altered primary cilia function to progressive kidney scarring in NPH remain poorly defined and therapeutic options allowing NPH patients to escape end-stage kidney disease are lacking.

Distinct proteins mutated in NPH interact with components of the Hippo pathway, an important regulator of cell fate. YAP (Yes-associated protein) overactivation has been shown to induce renal scarring while YAP inhibition showed protective effect in kidney diseases unrelated to NPH. Yet, the therapeutic potential of YAP inhibition in NPH has not been formerly assessed.

Here we studied the impact of both genetic and pharmacologic YAP inhibition on the NPH-like phenotype caused by a bi-allelic mutation of *Lkb1*, a ciliary kinase interacting with NPHP1. Contrary to non NPH renal disease, our results reveal an unexpected protective role of YAP in *Lkb1* mutant kidneys. Indeed, YAP genetic disruption drastically increase kidney disease burden in *Lkb1* deficient mice, while pharmacologic inhibition of YAP failed to improve their phenotype.

Collectively these results suggest that YAP inhibition is not a valid therapeutic strategy in NPH and suggest that LKB1 and YAP are parallel negative regulators of a yet uncharacterized pathway detrimental for kidney health.

## INTRODUCTION

Nephronophthisis (NPH) is an orphan genetic disease affecting the kidney. This recessive affection usually manifests with polyuria followed by a gradual reduction in kidney function related to progressive renal scarring. To date, no treatment is available for this affection, which is nonetheless the leading genetic cause of end-stage kidney disease in children^1,2^.

NPH is mostly caused by mutations affecting proteins that localize to primary cilia, solitary antenna-like organelles that protrude from the apical surface of most mammalian epithelial cells. Primary cilia emerged from the extension of tubulin doublets originating from the triplets forming the core of the mother centriole. Protein cargoes enter to and exit from the cilia through the transition zone, a complex protein sorting process taking place at the base of the cilium^3^. The most frequently mutated genes in NPH encode proteins that localize to the transition zone. Indeed, NPHP1, which mutations account for 25% of NPH cases, as well as NPHP4 and RPGRPI1L/NPHP8 are all core proteins of the transition zone^4^. Loss of function of *NPHP* genes does not impede ciliogenesis but perturbs cilia organization and/or signalling. Unfortunately, the dysregulated pathways responsible for kidney degeneration in NPH have not yet been solved, precluding the development of efficient therapies for the children and young adults affected by the disease.

One of the main difficulty faced studying the pathophysiologic events driving kidney damage in NPH is the overall lack of orthologous rodent models recapitulating all the disease features. Indeed, *Nphp1* inactivation does not lead to significant kidney fibrosis^5,6^ in mice and the same is true for the majority of NPH genes. One notable exception is *Glis2*, which inactivation consistently leads to kidney fibrosis in mice^7,8^, but accounts for only 0.1% of NPH cases in human.

Liver Kinase B1 (LKB1; encoded by *STK11*) is a ciliary kinase involved in the control of polarity and metabolism^9^. LKB1 interacts with NPHP1 and the bi-allelic disruption of *Stk11* in renal tubules of mice recapitulates the NPH phenotype^10^. Combining this model with the analysis of *Glis2* mutant mice and material derived from NPH patients, we further demonstrated that NPH is associated with a specific inflammatory signature that likely participates to kidney damage^11^. While NF-ΚB signalling has been identified as a contributor to kidney damage in *Glis2* deficient mice^12^, the molecular machinery driving kidney inflammation in NPH remains poorly defined.

The Hippo signalling pathway is an evolutionarily conserved kinase cascade that plays a fundamental role in several biologic processes such as embryonic development, organ size control, cell proliferation and apoptosis^13^. The main function of Hippo kinase is to phosphorylate the transcription co-activator YAP (Yes-associated protein) or its paralog TAZ/WWTR1 (Transcriptional coactivator with PDZ-binding domain). Unphosphorylated YAP and TAZ bind to transcriptional enhanced associate domain (TEAD1-4) transcription factors to regulate the expression of multiple genes in a cell and context specific fashion. Upon their phosphorylation by Hippo kinase, YAP and TAZ are targeted to degradation and/or sequestrated in the cytoplasm, shutting down the transcription of their target genes. In mammals, Hippo signalling consists in four serine/threonine kinases: the two upstream mammalian STE20-like protein kinases 1 and 2 (MST1/2; encoded by *STK4* and *STK3*, respectively) phosphorylate the effector large tumor suppressor kinases 1 and 2 (LATS1/2), which in turn phosphorylate YAP and TAZ causing their exclusion from the nuclear compartment^14,15^.

While Hippo pathway plays fundamental roles in kidney development^16,17^, several lines of evidence suggest that its deregulation drives kidney damage. Indeed, tubule specific inactivation of *Mst1* and *2* promotes renal inflammation and scarring largely through the induction of YAP target genes^18^. YAP and TAZ activation has been observed in autosomal dominant polycystic kidney disease (ADPKD)^19^ which is a genetic renal cystic disease caused by mutations affecting the ciliary proteins polycystins 1 and 2. In this context, the disruption of YAP and TAZ has been shown to reduce cystic disease burden in mice^20,21^. The idea that YAP/TAZ exerts pro-fibrotic function in the kidney is further supported by studies showing that verteporfin, a compound dissociating YAP/TEAD complex, reduces kidney fibrosis in genetic or acquired model of kidney diseases^22,23^.

Experimental data has linked NPH proteins to Hippo/YAP pathway. *In vitro*, NPHP4 binds and inhibits MST1^24^, while NPHP9/NEK8 has been shown to facilitate TAZ nuclear translocation^25^. On the other hand, NPHP9/NEK8 and NPHP16/ANKS6 pathogenic variants were reported to increase YAP transcriptional output in severe syndromic cystic dysplasia patients derived fibroblasts^26^ and late onset chronic kidney disease^27^. Verteporfin treatment rescues the developmental phenotype caused by *nek8* inhibition in zebrafish embryo^26^. In the same line, an unbiased siRNA screen identified LKB1 as a suppressor of YAP transcriptional activity^28^, while *Yap* inactivation has been shown to prevent liver growth caused by *Lkb1* disruption^28^.

Taken together these data suggest that NPH and Hippo/YAP signalling intersects, but the signification of this interaction for NPH kidney disease has not formerly been assessed. The growing list of drugs allowing the manipulation of Hippo and/or YAP activation prompted us to investigate this question.

## RESULTS

### Depletion of LKB1 or NPHP1 impairs Hippo signalling

To determine how LKB1 and NPHP1 intersect with Hippo pathway, we used cultured renal epithelial cells (MDCK) in which LKB1 or NPHP1 were depleted by shRNA-mediated knockdown^10^. Western blot experiments revealed that while nuclear YAP accumulated in *Lkb1* and *Nphp1* knockdown cells as compared to control cells (**Figure 1A**), cytoplasmic and nuclear TAZ levels were not modified. In parallel, the expression of *Ankrd1* (Ankyrin repeat domain 1), a prototypic YAP/TEAD transcriptional target, increased upon *Lkb1* or *Nphp1* shRNA-mediated knockdown (**Figure 1B**). To obtain further mechanistic insights, we examined the interaction between LKB1, NPHP1 and the upstream regulators of YAP nuclear localization. As previously described by others^28,29^, both LKB1 and NPHP1 co-immunoprecipitated with MST1 and LATS1 kinases *in vitro* (**Figure 1C**). Collectively, these data suggest that both LKB1 and NPHP1 interact with Hippo kinases to repress the nuclear translocation of YAP, thereby reducing the expression of YAP target genes.

**Figure 1.**
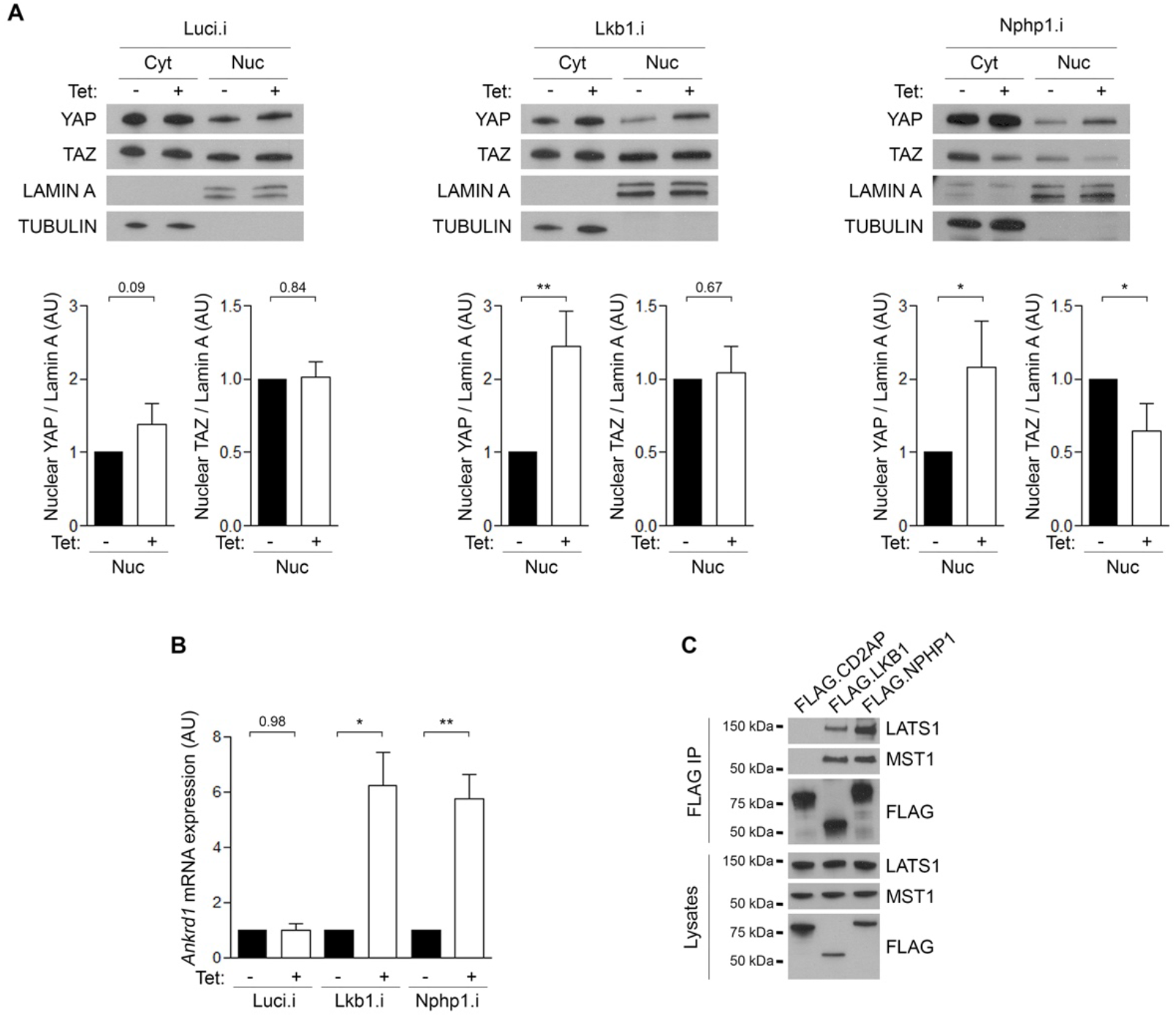
Depletion of LKB1 and NPHP1 impairs Hippo signaling. **(A)** Western blots and quantification of YAP/TAZ nucleo-cytoplasmic expression in MDCK cells expressing inducible shRNA against Luciferase (Luci.i), Lkb1 (Lkb1.i) and Nphp1 (Nphp1.i) 6 days after tetracycline induction (Tet). **(B)** Quantitative PCR of *Ankrd1* mRNA level in the same cell lines (n=5). **(C)** Immunoprecipitation (IP) from HEK 293T cells. Endogenous LATS1 and MST1 are enriched in the precipitates of FLAG.LKB1 and FLAG.NPHP1 but not FLAG.CD2AP. Representative Western blot of three independent experiments. Bars indicate mean ± SD. Paired t-test, * *P* < 0.05, ** *P* < 0.01. AU: arbitrary unit.

### Loss of *Lkb1* in the mouse kidney results in Hippo pathway dysregulation

In order to verify whether LKB1 could modulate Hippo pathway also *in vivo*, we took advantage of a mouse model characterized by *Lkb1* depletion specifically in the distal tubules (*Lkb1*^ΔTub^) causing a NPH-like phenotype^10^. Quantitative RT-PCR analysis revealed increased levels of known YAP target genes including *Cyr61* (cysteine-rich angiogenic inducer 61), *Ctgf* (connective tissue growth factor), *Ankrd1, Edn1* (endothelin 1) and *Areg* (amphiregulin) mRNA in the *Lkb1*^ΔTub^ mice compared to control mice as soon as 5 weeks after birth (**Figure 2A**). Moreover, the expression of these genes increased over time as higher levels were found in 23-week old *Lkb1*^ΔTub^ mice as compared to controls (**Figure 2A**). Immunohistochemistry revealed that YAP is strongly expressed in nuclei of *Lkb1*^ΔTub^ mice, especially in epithelial cells belonging to dilated tubules similarly to what was found in ADPKD^19^ (**Figure 2B**). Together, our *in vivo* and *in vitro* data revealed that LKB1 and NPHP1 promoted Hippo pathway through inhibition of YAP/TEAD transcriptional activity.

**Figure 2.**
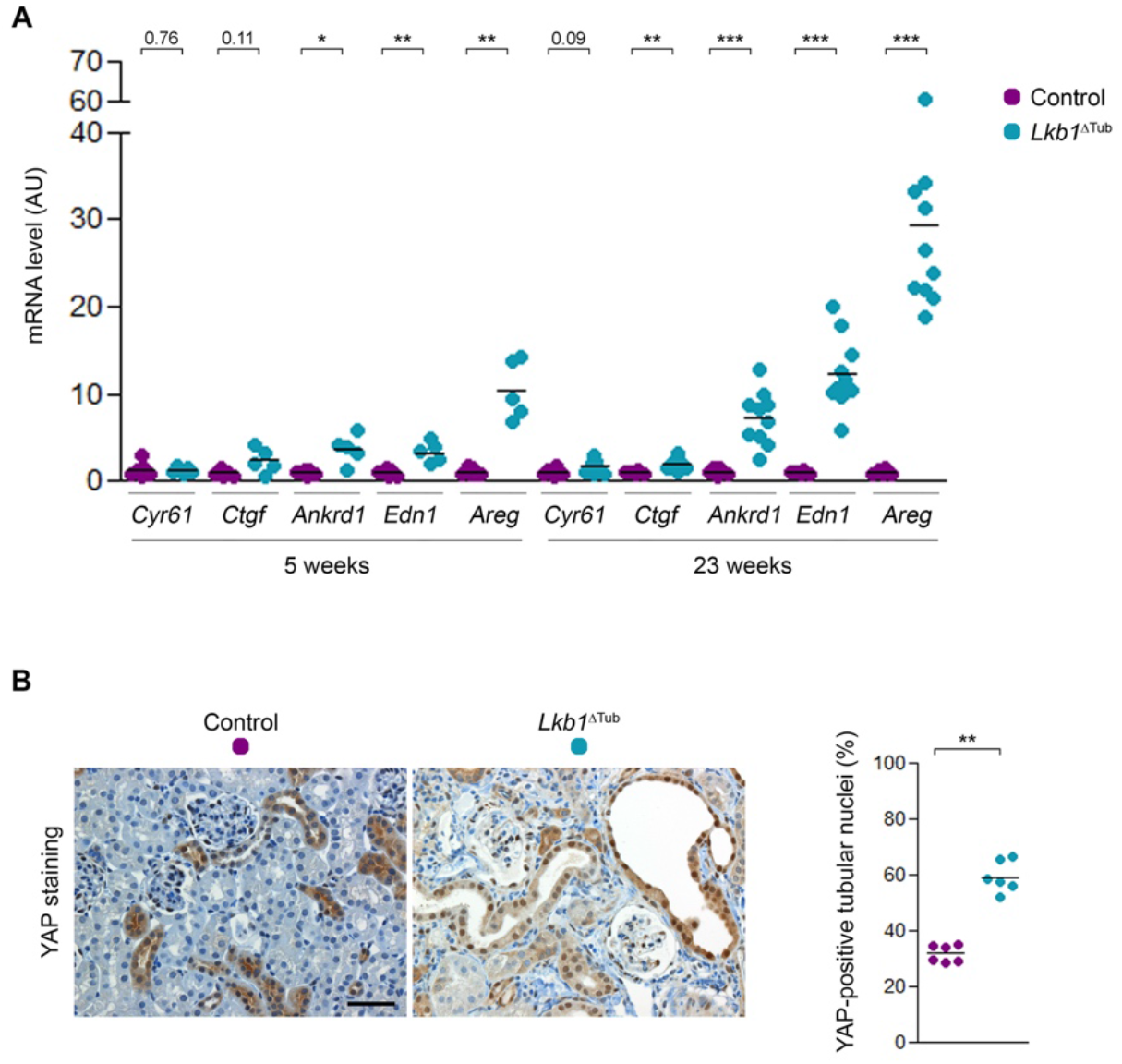
Loss of *Lkb1* in the mouse kidney results in Hippo pathway dysregulation and causes nuclear YAP imbalance. **(A)** Renal YAP transcriptional targets (*Cyr61, Ctgf, Ankrd1, Edn1, Areg*) mRNA content evaluated by quantitative PCR in control and *Lkb1*^ΔTub^ mice at 5 and 23 weeks of age. Each dot represents one individual. Bars indicate mean. Mann-Whitney t-test, * *P* < 0.05, ** *P* < 0.01, *** *P* < 0.001. AU: arbitrary unit. **(B)** YAP immunostaining and quantification in kidneys from control and *Lkb1*^ΔTub^ mice at 23 weeks. Representative images of n=5 mice/group. Scale bar: 50 µm.

### Loss of tubular *Yap* causes premature death, urine concentration defect and rapid renal function decline in *Lkb1*^ΔTub^ mice

To determine if the deletion of *Yap* together with *Lkb1* would prevent the NPH-like renal disease, we produced an allelic series by crossing *Lkb1*^ΔTub^ mice with mice bearing *Yap* floxed alleles and compared them with littermate controls. At 4 weeks of age, kidneys of *Lkb1*^ΔTub^ mice displayed reduced LKB1 expression measured by RT-PCR and immunohistochemistry (**Supplementary Figure 1A-B**). Similarly, *Yap* mRNA and protein expression were decreased in *Yap*^ΔTub^ kidneys and associated with enhanced *Cyr61* mRNA expression without affecting *Taz* mRNA level (**Supplementary Figure 1C-F**). Macroscopic inspection of the kidneys from control and mutated mice revealed that *Yap*^ΔTub^ kidneys resemble the control ones, while *Lkb1*^ΔTub^ kidneys showed a discrete surface granular irregularities, as previously described^10^. Surprisingly, kidneys carrying a bi-allelic inactivation of *Yap* and *Lkb1* showed drastic parenchymal thinning with medulla atrophy resulting in a balloon-like aspect (**Figure 3A**). Importantly, renal pelvis from double mutant animals were not distended excluding ureteral obstruction. As a consequence *Lkb1*^ΔTub^; *Yap*^ΔTub^ mice displayed smaller kidneys compared to controls (**Figure 3B** and **Supplementary Figure 2A-B**). Contrary to other animals, *Lkb1*^ΔTub^; *Yap*^ΔTub^ mice did not survive for more that 8 weeks (**Figure 3C**). As soon as 4 weeks of age, all the animals carrying a bi-allelic inactivation of either *Lkb1* or *Yap* displayed a dramatic urine concentration defect as compared to littermate controls (**Figure 3 D**). Remarkably, only *Lkb1*^ΔTub^ mice with allelic dosage of *Yap* inactivation showed a reduced kidney function proportional to *Yap* inactivation (**Figure 3E**). By contrast, *Yap*^ΔTub^ mice showed a normal kidney function. Collectively, these results demonstrate that the combination of *Lkb1* and *Yap* inactivation in distal tubules worsen the *Lkb1*^ΔTub^ phenotype.

**Figure 3.**
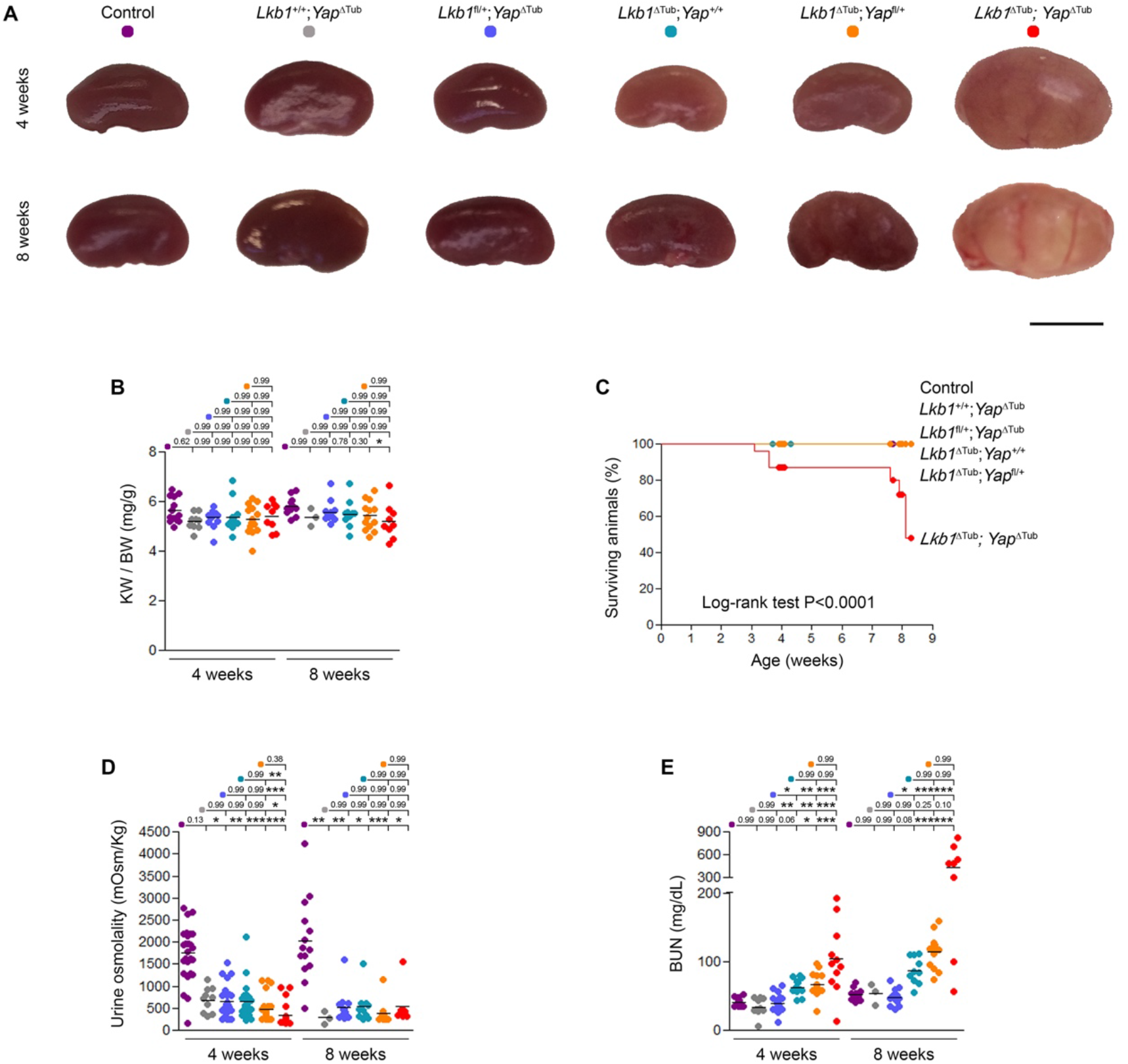
Loss of tubular *Yap* causes premature death and rapid renal function decline in *Lkb1*^ΔTub^ mice. **(A)** Representative kidneys from control, *Lkb1*^+/+^; *Yap*^ΔTub^, *Lkb1*^fl/+^; *Yap*^ΔTub^, *Lkb1*^ΔTub^; *Yap*^+/+^, *Lkb1*^ΔTub^; *Yap*^fl/+^ and *Lkb1*^ΔTub^; *Yap*^ΔTub^ mice at 4 and 8 weeks. Scale bar: 5 mm. **(B)** Kidney weight (KW) to body weight (BW) ratio in the 6 groups of animals at 4 and 8 weeks. **(C)** Kaplan–Meier survival curves of the same groups of animals. **(D-E)** Urine osmolality (D), plasma blood urea nitrogen (BUN, E) in the 6 groups of animals at 4 and 8 weeks. Each dot represents one individual mouse. Bars indicate mean. Kruskal-Wallis test, * *P* < 0.05, ** *P* < 0.01, *** *P* < 0.001. AU: arbitrary unit.

### Genetic inhibition of tubular *Yap* promotes tubular lesions and fibrosis in *Lkb1* mutant mice

Thickened tubular basement membranes, tubular dilatation and atrophy, interstitial cell infiltration and renal fibrosis are the main features that characterize *Lkb1*^ΔTub^ kidneys. Renal histology revealed that all these features were exacerbated in *Lkb1*^ΔTub^ in proportion to the reduction of *Yap* gene dosage (**Figure 4A-B** and **Supplementary Figure 2**). Consistently, *Yap* deletion in *Lkb1*^ΔTub^ kidneys proportionally increased the mRNA expression of two tubular injury markers, *Lcn2* and *Kim1* (**Figure 4C-D**). Sirius red staining confirmed the increase in renal fibrosis in *Lkb1*^ΔTub^ kidneys, which was more pronounced in *Lkb1*^ΔTub^; *Yap*^ΔTub^ kidneys. Markers of interstitial fibrosis, such as collagens or pro-fibrotic cytokine levels (*Tgfb1, Pdgfb*) were increased proportionally to *Yap* gene dosage reduction (**Figure 4E-G** and **Supplementary Figure 2**). By contrast, *Yap* inactivation in wild-type *Lkb1* animals (*Yap*^ΔTub^) did not lead to kidney damage at least in this time frame. Overall, our data indicate that the mid-term maintenance of *Lkb1* deficient kidney structure and function is critically dependent of YAP.

**Figure 4.**
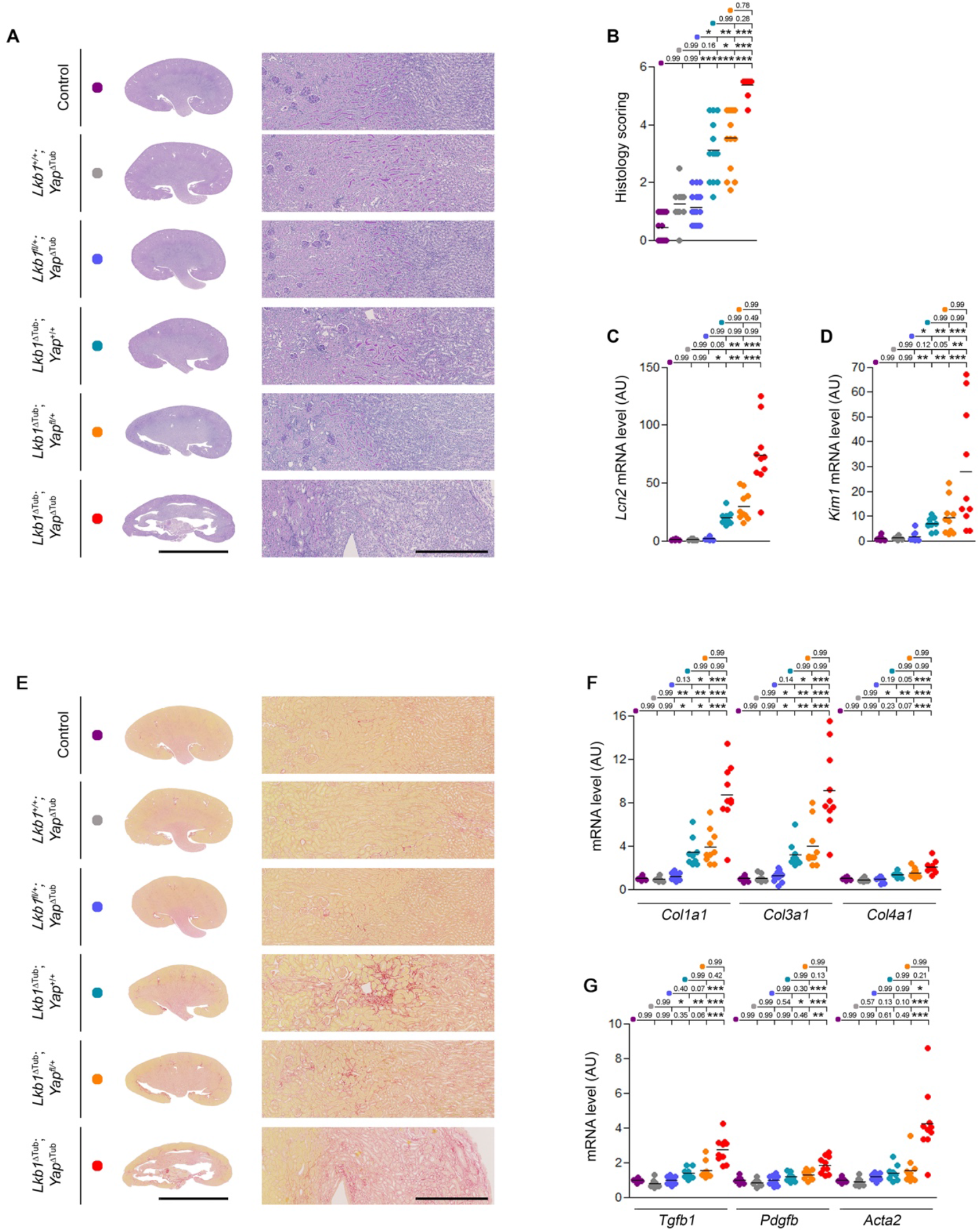
*Yap* ablation enhances tubular lesions and fibrosis in *Lkb1* mutant mice. **(A)** Representative PAS stained kidney sections from control, *Lkb1*^+/+^; *Yap*^ΔTub^, *Lkb1*^fl/+^; *Yap*^ΔTub^, *Lkb1*^ΔTub^; *Yap*^+/+^, *Lkb1*^ΔTub^; *Yap*^fl/+^ and *Lkb1*^ΔTub^; *Yap*^ΔTub^ mice at 4 weeks. Scale bars: 5mm (left panel), 500µm (right panel). **(B)** Histology lesion scoring in kidneys from the same groups of animals. **(C-D)** *Lcn2* (C) and *Kim1* (D) renal mRNA expression at 4 weeks. **(E)** Representative sirius red stained kidney sections of the same groups of animals at 4 weeks. Scale bars: 5mm (left panel), 500µm (right panel). **(F)** Renal collagen mRNA content evaluated by quantitative PCR at 4 weeks. **(G)** *Tgfb1, Pdgfb* and *Acta2* mRNA expression in kidneys from the same mice at 4 weeks. **(B-D, F-G)** Each dot represents one individual mouse. Bars indicate mean. Kruskal-Wallis test, * *P* < 0.05, ** *P* < 0.01, *** *P* < 0.001. AU: arbitrary unit.

### Tubular *Yap* inactivation promotes renal inflammation in *Lkb1*^ΔTub^ mice

We previously showed that NPH is characterized by a complex and specific cytokine signature that is associated with an early kidney infiltration by macrophages, neutrophils and T cells^11^. Thus, we evaluated the inflammatory status of *Lkb1*^ΔTub^ mice when *Yap* was inactivated. Quantitative RT-PCR analysis revealed a global upregulation of the pro-inflammatory cytokines identified in NPH that directly correlated with the level of *Yap* deletion (**Figure 5A** and **Supplementary Figure 3**). As these secreted factors are involved in the recruitment of different immune cell populations, we then performed immunohistochemistry staining for macrophages (F4/80), T cells (CD3) and neutrophils (Ly6B.2). Supporting the molecular results, *Lkb1*^ΔTub^; *Yap*^ΔTub^ kidneys presented increased kidney infiltration by macrophages, T cells and neutrophils (**Figure 5B-F** and **Supplementary Figure 4**). Of note, *Yap* inactivation did not result in renal inflammation in animal carrying at least one functional copy of *Lkb1* (*Yap*^ΔTub^ and *Lkb1*^fl/+^; *Yap*^ΔTub^). These results confirmed that the specific inflammatory signature previously identified is associated with the development of the disease and indicated that tubular YAP limits kidney inflammation in NPH.

**Figure 5.**
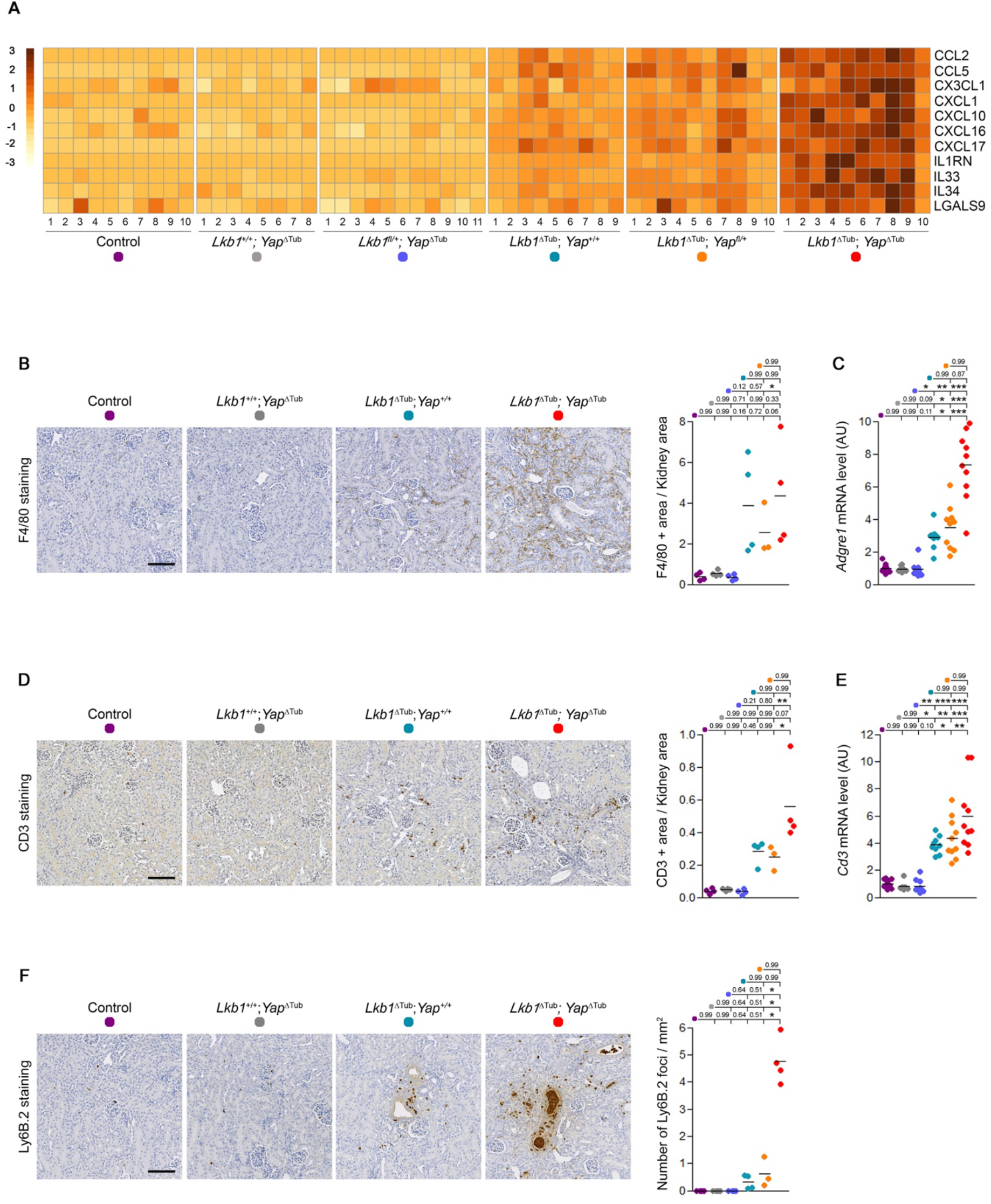
Tubular *Yap* inactivation promotes renal inflammation in *Lkb1*^ΔTub^ mice. **(A)** Heatmap showing Z-scores computed on mRNA expression measured by quantitative PCR of the indicated cytokines in kidneys from 4 weeks old control, *Lkb1*^+/+^; *Yap*^ΔTub^, *Lkb1*^fl/+^; *Yap*^ΔTub^, *Lkb1*^ΔTub^; *Yap*^+/+^, *Lkb1*^ΔTub^; *Yap*^fl/+^ and *Lkb1*^ΔTub^; *Yap*^ΔTub^ mice. See also **Supplementary Figure 3. (B)** Representative images and quantification of F4/80 (macrophages) immunostaining of kidney sections from 4 weeks old animals. Scale bar: 100µm. **(C)** *Adgre1* mRNA expression in the same mice. **(D)** Representative images and quantification of CD3 (T cells) immunostaining of kidney sections from 4 weeks old mice. Scale bar: 100µm. **(E)** *Cd3* mRNA expression in kidneys from the same animals. **(F)** Representative images and quantification of Ly-6B.2 (neutrophils) immunostaining of kidney sections from 4 weeks old mice. Scale bar: 100µm. **(B-F)** Each dot represents one individual mouse. Bars indicate mean. Kruskal-Wallis test, * *P* < 0.05, ** *P* < 0.01, *** *P* < 0.001. AU: arbitrary unit.

### Pharmacological inhibition of YAP enhances renal inflammation in *Lkb1* mutant mice

YAP can be targeted pharmacologically by verteporfin, which inhibits the formation of the transcriptomic complex YAP-TEAD. We treated control and *Lkb1*^ΔTub^ mice with verteporfin (*i*.*p*. 100mg/kg) from 8 to 12 weeks of age. While verteporfin treatment had no impact on urine concentration defect or kidney function in *Lkb1*^ΔTub^ mice, we observed some lethality and higher mRNA expression of *Lcn2* tubular injury marker in verteporfin-treated *Lkb1*^ΔTub^ animals (**Figure 6A-B** and **Supplementary 5A-G**). Verteporfin treatment did not reduce interstitial fibrosis nor fibrotic genes expression in *Lkb1*^ΔTub^ mice (**Supplementary Figure 5H-J**). However, verteporfin treatment in *Lkb1*^ΔTub^ animals tends to increase the expression of some inflammatory cytokines and exarcerbate infiltration of the kidney by immune cells (**Figure 6D-Q**). Overall, our genetic and pharmacologic inhibition of YAP demonstrated that reducing YAP gene dosage or YAP/TEAD transcriptional activity is not a viable strategy to modulate NPH progression.

**Figure 6.**
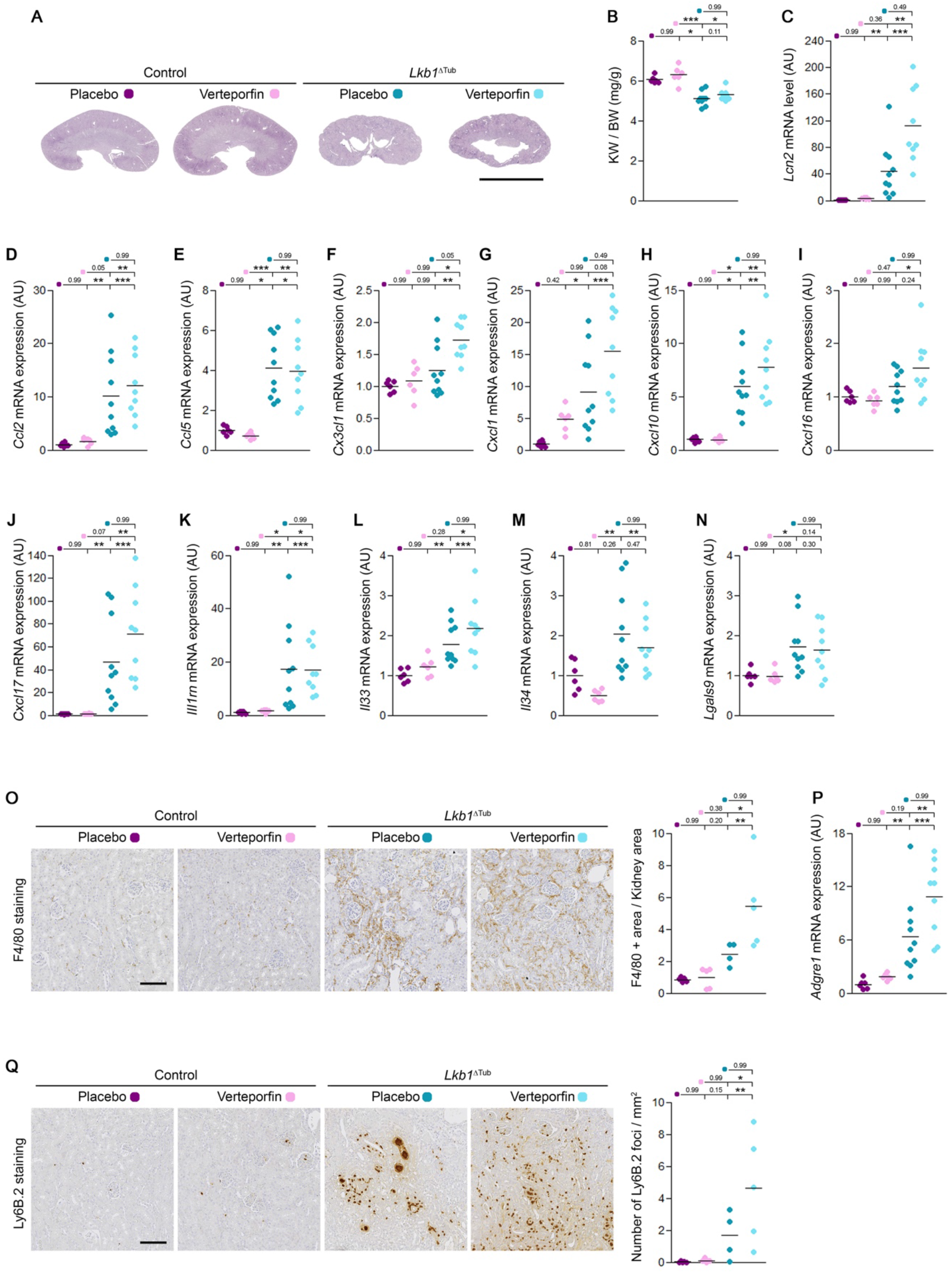
Pharmacological inhibition of YAP enhances renal inflammation in *Lkb1* mutant mice. **(A)** Representative PAS stained kidney sections from 12 weeks-old control and *Lkb1*^ΔTub^ mice administered *i*.*p*. with vehicle (DMSO) or verteporfin (100mg/Kg) every other day for 1 month. Scale bars: 5mm. See also **Supplementary Figure 5. (B)** Kidney weight (KW) to body weight (BW) ratio of the same groups of mice at 12 weeks. **(C-N)** Quantification of *Lcn2* (C), *Ccl2* (D), *Ccl5* (E), *Cx3cl1* (F), *Cxcl1* (G), *Cxcl10* (H), *Cxcl16* (I), *Cxcl17* (J), *Il1rn* (K), *Il33* (L), *Il34* (M) and *Lgals9* (N) mRNA abundance in kidneys from 12 weeks old control and *Lkb1*^ΔTub^ mice treated or not with verteporfin. **(O)** Representative images and quantification of F4/80 (macrophages) immunostaining of kidney sections from 12 weeks old animals. Scale bar: 100µm. **(P)** *Adgre1* mRNA expression in the same mice. **(Q)** Representative images and quantification of Ly-6B.2 (neutrophils) immunostaining of kidney sections from 12 weeks old mice. Scale bar: 100µm. **(B-Q)** Each dot represents one individual mouse. Bars indicate mean. Kruskal-Wallis test, * *P* < 0.05, ** *P* < 0.01, *** *P* < 0.001. AU: arbitrary unit.

## DISCUSSION

The aim of this study was to determine if YAP inhibition could represent a valid therapeutic target in NPH using *Lkb1*^ΔTub^ mouse model. Our results clearly demonstrate that, in this model, YAP activation does not contribute to kidney lesions. On the opposite, we observed that tubule specific *Yap* inactivation drastically worsened kidney damage associated with *Lkb1* loss of function. Similarly, treating mice with the systemic YAP/TEAD inhibitor verteporfin did not slow the progression of NPH-like renal disease and even tends to worsen its inflammatory aspect.

As we used a non-orthologous NPH model of the disease that recapitulates the fibrotic feature observed in human bearing bi-allelic *NPHP1* deletion, we cannot rule out that YAP inhibition may exert some positive effects on the less frequent forms of NPH characterized by a cystic phenotype reminiscent of ADPKD. In this regard, it would be interesting to perform similar studies in *jck* mice, which bear a spontaneous mutation in *Nphp9/Nek8* that cause cystogenesis associated with YAP activation^26^.

Interestingly, while YAP genetic disruption drastically worsened the kidney phenotype of *Lkb1*^ΔTub^ mice, verteporfin treatment was mostly neutral. This discrepancy may reflect an incomplete inhibition of YAP transcriptional activity by verteporfin. Alternatively, this observation could also pinpoint a TEAD independent functions of YAP in *Lkb1*^ΔTub^ kidneys. Besides its function as a co-activator of TEAD, YAP has been shown to bind other transcription factors such as RUNX2/3, p73, SMADs, ERBB^30^. In addition, YAP also modulates important signaling pathways independently of transcription factors. Such regulation has been notably demonstrated for important pro-inflammatory pathways such as NF-κB, which has also been implicated in the kidney disease caused by *Glis2* inactivation^12^.

While our results do not support YAP inhibition as a promising therapy for NPH, they unveil an unforeseen protective role of YAP in this disease. Our results suggest a model in which LKB1 and YAP are parallel negative regulators of a molecular pathway responsible for the development of kidney lesions in NPH. Further experiments are needed to validate this model and may lead to the identification of molecular events responsible for NPH.

## MATERIALS and METHODS

### Mice

Mice were housed in a specific pathogen-free facility, fed *ad libitum* and housed at constant ambient temperature in a 12-hour day/night cycle. Breeding and genotyping were done according to standard procedures.

*Lkb1*^ΔTub^ mice were previously described^10^. *Yap*^flox/flox^ mice (C57BL/6J) were kindly provided by Prof. Eric Olson (University of Texas South western Medical Center, Dallas, USA)^31^ and were backcrossed for 2 generations with *Lkb1*^ΔTub^ mice. The progeny was then intercrossed to generate mice with allelic dosage of *Lkb1* into tubule-specific *Yap* knockout: *Yap* knockout with wild-type *Lkb1* (further referred to as *Lkb1*^+/+^; *Yap*^ΔTub^) or heterozygous floxed *Lkb1* allele (further referred to as *Lkb1*^fl/+^; *Yap*^ΔTub^); and mice with allelic dosage of *Yap* into tubule-specific *Lkb1* knockout: *Lkb1* knockout with wild-type *Yap* (further referred to as *Lkb1*^ΔTub^; *Yap*^+/+^), heterozygous *Yap* (further referred to as *Lkb1*^ΔTub^; *Yap*^fl/+^) or homozygous floxed *Yap* alleles (further referred to as *Lkb1*^ΔTub^; *Yap*^ΔTub^). Littermates lacking *KspCre* transgene were used as controls. Experiments were conducted on both females and males.

For Verteporfin experiment, only control and *Lkb1*^ΔTub^ male mice were used. Verteporfin (Sigma, SML0534) was dissolved in DMSO (100 mg/mL), aliquoted, and stored at −80°C. Working solution was prepared at 10 mg/mL in PBS freshly before use. Eight-week old mice were administered *i*.*p*. with vehicle or verteporfin solution at a dose of 100 mg/kg every other day for 1 month. Animals were sacrificed 4 hours after the last injection.

### Urine and Plasma Analyses

Spot urine samples were obtained from mice the day of sacrifice. Urine osmolality was measured with a freezing point depression osmometer (Micro-Osmometer from Knauer). For Verteporfin protocol, 8-hour urine samples were obtained from mice housed in individual boxes without access to water and food. Urine excretion was measured.

Retro-orbital blood was collected from anesthetized mice. Plasma blood urea nitrogen (BUN) was measured using a Konelab 20i Analyzer (Thermo Scientific).

### Morphological Analysis

Mouse kidneys were fixed in 4% paraformaldehyde, embedded in paraffin, and 4µm sections were stained with periodic acid-Schiff (PAS) or Picrosirius Red. Stained full size images were recorded using a whole slide scanner Nanozoomer 2.0 (Hamamatsu) equipped with a 20x/0.75 NA objective coupled to NDPview software (Hamamatsu). The severity of renal pathology was graded in a blinded fashion by two independent observers assessing the overall lesions on the whole kidney section stained with PAS. A 7-point scale was defined, in which 0 indicated normal kidney architecture, 3 indicated tubular atrophy, tubular basement thickening and cell infiltration and 6 characterized by loss of medulla and complete disorganization of the remaining renal parenchyma. For verteporfin experiment, only four scores were defined ranging from score 0 normal kidney architecture to score 3 associating tubular atrophy, tubular basement thickening and cell infiltration.

### Cell culture

Madin-Darby canine kidney cells (MDCK, kind gift from Prof. Kai Simons, MPI-CBG, Dresden, Germany) were cultured using Dulbecco’s modified Eagle’s medium supplemented with 10% fetal bovine serum and 1% penicillin–streptomycin (Sigma). Generation of MDCK cell lines for tetracycline-inducible knockdown of target genes was previously described^10^. The shRNA targeting sequences used were as follows: Lkb1-i (5’-GCTGGTGGACGTGTTATAC-3’), Luciferase (5’-CGTACGCGGAATACTTCGA-3’) and Nphp1-i (5’-GGTTCTCAGTAGACATGTA-3’). For nucleo-cytoplasmic experiment, 150,000 cells/cm^2^ were seeded with or without tetracycline (5 µg/ml) for 6 days on 6 well plate (Greiner, 657 160). Cells were lysed in Pierce NE-PER^™^ buffer (Thermo Fisher Scientific) to extract both nuclear and cytoplasmic proteins. In parallel, 150,000 cells/cm^2^ were seeded with or without tetracycline (5 µg/mL) for 6 days on polycarbonate membranes (COSTAR, 3401). Total RNAs were extracted after 5 hours of serum deprivation and qRT–PCR was performed.

Human embryonic kidney (HEK 293T) cells (ATCC Promocell; CRL-11268) were cultured using Dulbecco’s modified Eagle’s medium supplemented with 10% fetal bovine serum (Biochrom). Plasmids were transiently transfected using calcium. Tagged constructs for CD2AP, NPHP1, and LKB1 were previously described^4^. Cells were lysed in IP buffer (20 mM Tris-HCl pH 7.5, 50mM NaCl, 1% Triton X100, 15mM Na_4_P_2_O_7_ and 0.1mM EDTA) supplemented with 2 mM Na_3_VO_4_, 50 mM NaF, 5 mM b-glycerophosphate, and protease inhibitor cocktail tablet (Roche) using a 20-G needle. Equal amounts of proteins were incubated with anti-FLAG M2 affinity gel (Sigma Aldrich, A2220) and processed by Western blotting.

All cells were regularly tested for mycoplasma contamination and were mycoplasma-free.

### Quantitative RT-PCR

Total RNAs were obtained from cells or mouse kidneys using RNeasy Mini Kit (Qiagen) and reverse transcribed using High Capacity cDNA Reverse Transcription Kit (Applied Biosystems) according to the manufacturer’s protocol. Quantitative PCR were performed with iTaq™ Universal SYBR® Green Supermix (Bio-Rad) on a CFX384 C1000 Touch (Bio-Rad). *Hprt, Ppia and Rpl13* were used as normalization controls^32^. Each biological replicate was measured in technical duplicates. The primers used for qRT-PCR are listed in Supplementary Table 1.

Heatmap was generated using R v4.1.0 and the R package pheatmap v1.0.12 (note: heatmaps displays Z-scores computed on the expression levels of the identified cytokines, measured by qPCR).

### Western blot

Cells were lysed in IP buffer or Pierce NE-PER^™^ buffer (Thermo Fisher Scientific) using a 20-G needle. Protein content was determined with Pierce BCA protein assay kit (Thermo Fisher Scientific). Equal amounts of proteins were resolved on 4–15% Mini-PROTEAN® TGX^™^ gel (Bio-Rad) under reducing conditions, transferred, incubated with primary and secondary antibodies, and visualized on film according to standard protocols. Band density was calculated and normalized using LabImage 1D L340 software. Antibodies used are listed in Supplementary Table 2.

### Immunohistochemistry

Four-micrometer sections of paraffin-embedded kidneys were submitted to antigen retrieval and avidin/biotin blocking (Vector, SP-2001). Sections were incubated with primary antibody followed by biotinylated antibody, HRP-labeled streptavidin (Southern Biotech, 7100-05, 1:500), and 3,30-diaminobenzidine-tetrahydrochloride (DAB) revelation. Antibodies used are listed in Supplementary Table 2. Full size images were recorded using a whole slide scanner Nanozoomer 2.0 (Hamamatsu) coupled to NDPview software. For macrophage quantification, stained area was measured with ImageJ software from full sized kidney images and visualized as the ratio of stained DAB surface to total kidney section area. For neutrophil quantification, we counted manually the number of foci in the whole kidney section. Foci were defined as 4 or more neutrophils surrounding a tubule. For T cell quantification, 10 randomly selected fields of view were measured with ImageJ software. For all quantification, glomerular and intra-tubular positive stainings were removed from the analysis.

### Statistical analysis

Data were expressed as means. Differences between groups were evaluated using Mann-Whitney test when only two groups were compared. When testing more comparisons, Kruskal-Wallis test was used. The statistical analysis was performed using GraphPad Prism V8 software. All image analyses and mouse phenotypic analyses were performed in a blinded fashion.

### Study approval

All animal experiments were conducted according to the guidelines of the National Institutes of Health *Guide for the Care and Use of Laboratory Animals*, as well as the German and French laws for animal welfare, and were approved by regional authorities (Regierungspräsidium Freiburg G-13/18 and Ministère de l’Enseignement, de la Recherche et de l’Innovation APAFIS#2020051216078531).

## ACKNOWLEDGEMENTS

We are grateful to Prof. Eric Olson for sharing *Yap*^fl/fl^ mice, and to Dr Frank Bienaimé for critically reading the manuscript. We thank the members of the mouse histology and breeding facilities (S.F.R Necker INSERM US24, Paris, France), and of the mouse renal physiology facility (CRC, Paris, France) for technical assistance.

Marceau Quatredeniers and Amandine Viau were supported by a public grant “RHU-C’IL-LICO” overseen by the Agence Nationale de la Recherche (grant number: ANR-17-RHUS-0002), E. Wolfgang Kuehn was supported by Deutsche Forschungsgemeinschaft KU1504/7-1 and KU1504/8-1, Sophie Saunier was supported by the Institut National de la Santé et de la Recherche Médicale (INSERM), the Ministère de l’Education Nationale, de la Recherche et de la Technologie (MRT), by a State funding from the Agence Nationale de la Recherche (grant references: ANR-10-IAHU-01, ANR-17-RHUS-0002).

