## Supplementary Material for "YAP restricts renal inflammation and mitigates kidney damage in nephronothisis related kidney disease"

**Supplementary Data**

**Table of contents**

**Supplementary Figure Legends**

**Supplementary Figure 1.** Validation of tubular *Lkb1* and *Yap* depletion in mice.

**Supplementary Figure 2.** Characterization of renal lesions after loss of tubular *Yap* in *Lkb1* mutant mice at 8 weeks.

**Supplementary Figure 3.** Pro-inflammatory cytokines are enhanced after loss of tubular *Yap* in *Lkb1*$^{\Delta Tub}$ mice.

**Supplementary Figure 4.** Characterization of renal inflammation after *Yap* depletion in *Lkb1* mutant mice at 8 weeks.

**Supplementary Figure 5.** Characterization of pharmacological YAP inhibition in *Lkb1*$^{\Delta Tub}$ mice.

**Supplementary Tables**

**Supplementary Table 1**: Primer pairs used for qRT-PCR.

**Supplementary Table 2**: Antibodies list.

### Supplementary Figure 1

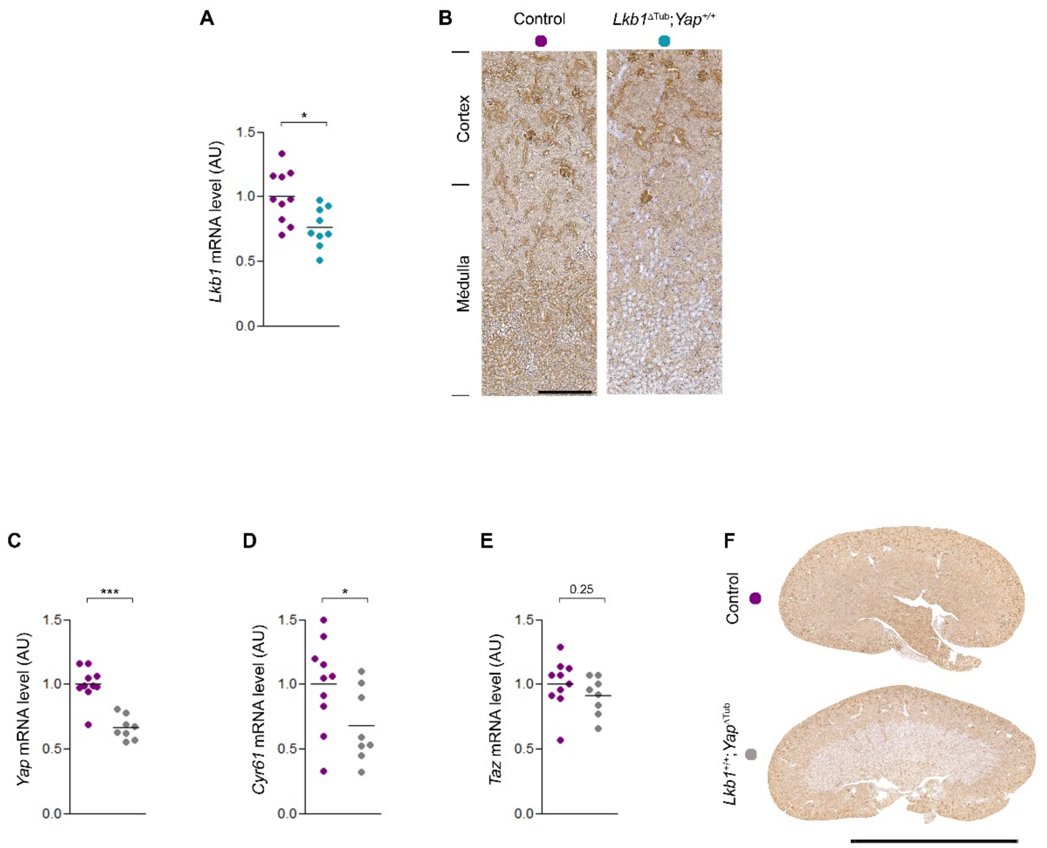


**Supplementary Figure 1. Validation of tubular *Lkb1* and *Yap* depletion in mice.**

**(A)** *Lkb1* mRNA expression in kidneys from 4 week old control and *Lkb1*$^{\Delta Tub}$ mice. **(B)** LKB1 immunostaining in kidneys from control and *Lkb1*$^{\Delta Tub}$ mice at 4 weeks. Representative images of n=5/group. Scale bar: 250µm. **(C-E)** *Yap* (C), *Cyr61* (D) and *Taz* (E) mRNA expression in kidneys from 4 week old control and *Yap*$^{\Delta Tub}$ mice. **(F)** YAP immunostaining in kidneys from control and *Lkb1*$^{\Delta Tub}$ mice at 4 weeks. Representative images of n=5/group. Scale bar: 5mm. **(A, C-E)** Each dot represents one individual mouse. Bars indicate mean. Mann-Whitney test, * $P$ < 0.05, *** $P$ < 0.001. AU: arbitrary unit.

**Supplementary Figure 2**

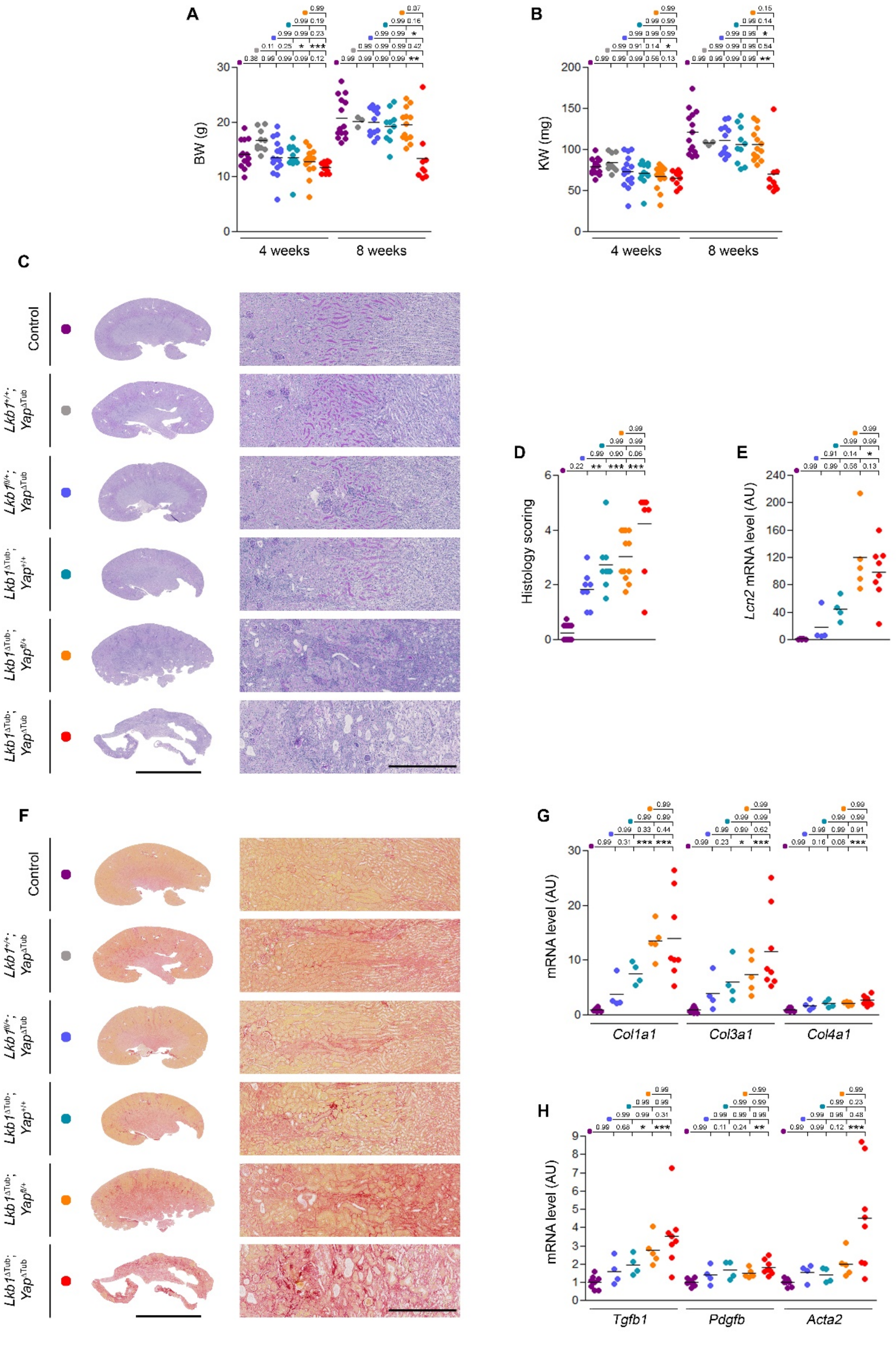


**Supplementary Figure 2. Characterization of renal lesions after loss of tubular *Yap* in *Lkb1* mutant mice at 8 weeks.**

**(A)** Body weight (BW) of control, *Lkb1*$^{+/+}$; *Yap*$^{\Delta Tub}$, *Lkb1*$^{fl/+}$; *Yap*$^{\Delta Tub}$, *Lkb1*$^{\Delta Tub}$; *Yap*$^{+/+}$, *Lkb1*$^{\Delta Tub}$; *Yap*$^{fl/+}$ and *Lkb1*$^{\Delta Tub}$; *Yap*$^{\Delta Tub}$ mice at 4 and 8 weeks. **(B)** Kidney weight (KW) of the same groups of mice at 4 and 8 weeks. **(C)** Representative PAS stained kidney sections at 8 weeks. Scale bars: 5mm (left panel), 500µm (right panel). **(D)** Histology lesion scoring in kidneys from the same animals at 8 weeks. **(E)** *Lcn2* renal mRNA expression at 8 weeks. **(F)** Representative sirius red stained kidney sections at 8 weeks. Scale bars: 5mm (left panel), 500µm (right panel). **(G)** Renal collagen mRNA content evaluated by quantitative PCR at 8 weeks. **(H)** *Tgfb1*, *Pdgfb* and *Acta2* mRNA expression in kidneys from the same mice at 8 weeks. **(A-B, D-E, G-H)** Each dot represents one individual mouse. Bars indicate mean. Kruskal-Wallis test, * $P < 0.05$, ** $P < 0.01$, *** $P < 0.001$. AU: arbitrary unit.

**Supplementary Figure 3**

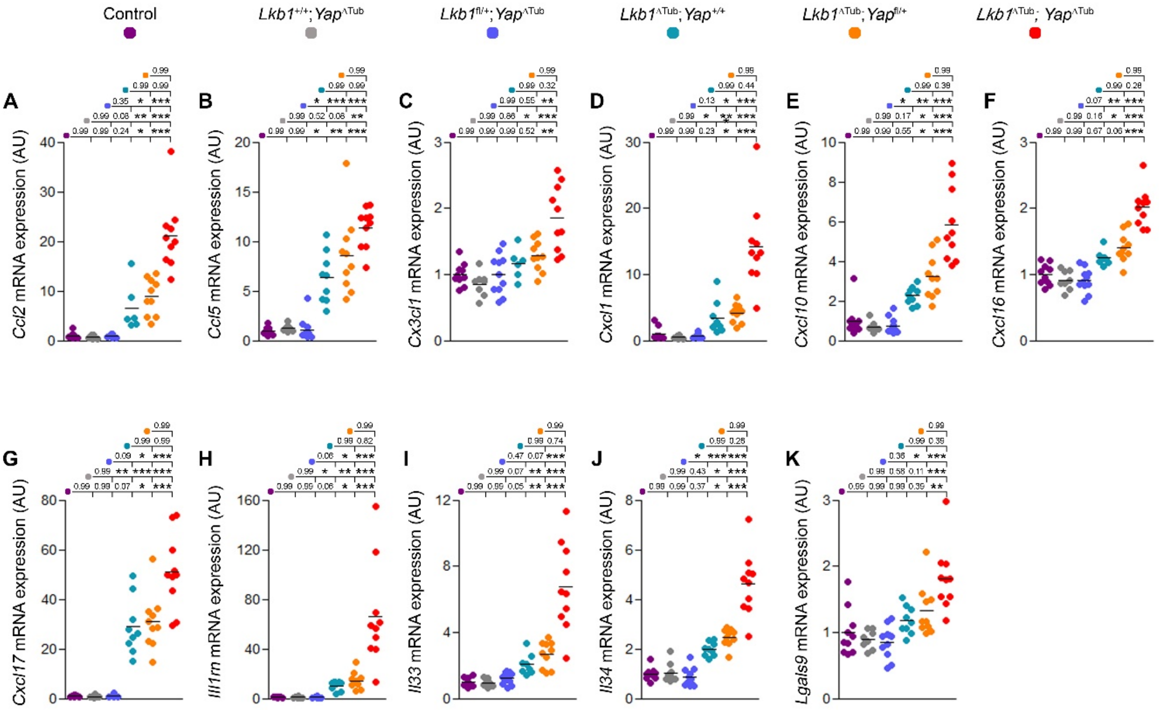


**Supplementary Figure 3. Pro-inflammatory cytokines are enhanced after loss of tubular *Yap* in *Lkb1*$^{\Delta Tub}$ mice.**

**(A-K)** Quantification of *Ccl2* (A), *Ccl5 (*B), *Cx3cl1* (C), *Cxcl1* (D), *Cxcl10* (E), *Cxcl16* (F), *Cxcl17* (G), *Il1rn* (H), *Il33* (I), *Il34* (J) and *Lgals9* (K) mRNA abundance in kidneys from 4 week old control, *Lkb1*$^{+/+}$; *Yap*$^{\Delta Tub}$, *Lkb1*$^{fl/+}$; *Yap*$^{\Delta Tub}$, *Lkb1*$^{\Delta Tub}$; *Yap*$^{+/+}$, *Lkb1*$^{\Delta Tub}$; *Yap*$^{fl/+}$ and *Lkb1*$^{\Delta Tub}$; *Yap*$^{\Delta Tub}$ mice. Each dot represents one individual mouse. Bars indicate mean. Kruskal-Wallis test, * $P < 0.05$, ** $P < 0.01$, *** $P < 0.001$. AU: arbitrary unit. A heatmap showing Z-scores computed on these mRNA expression is shown in Figure 5.

### Supplementary Figure 4

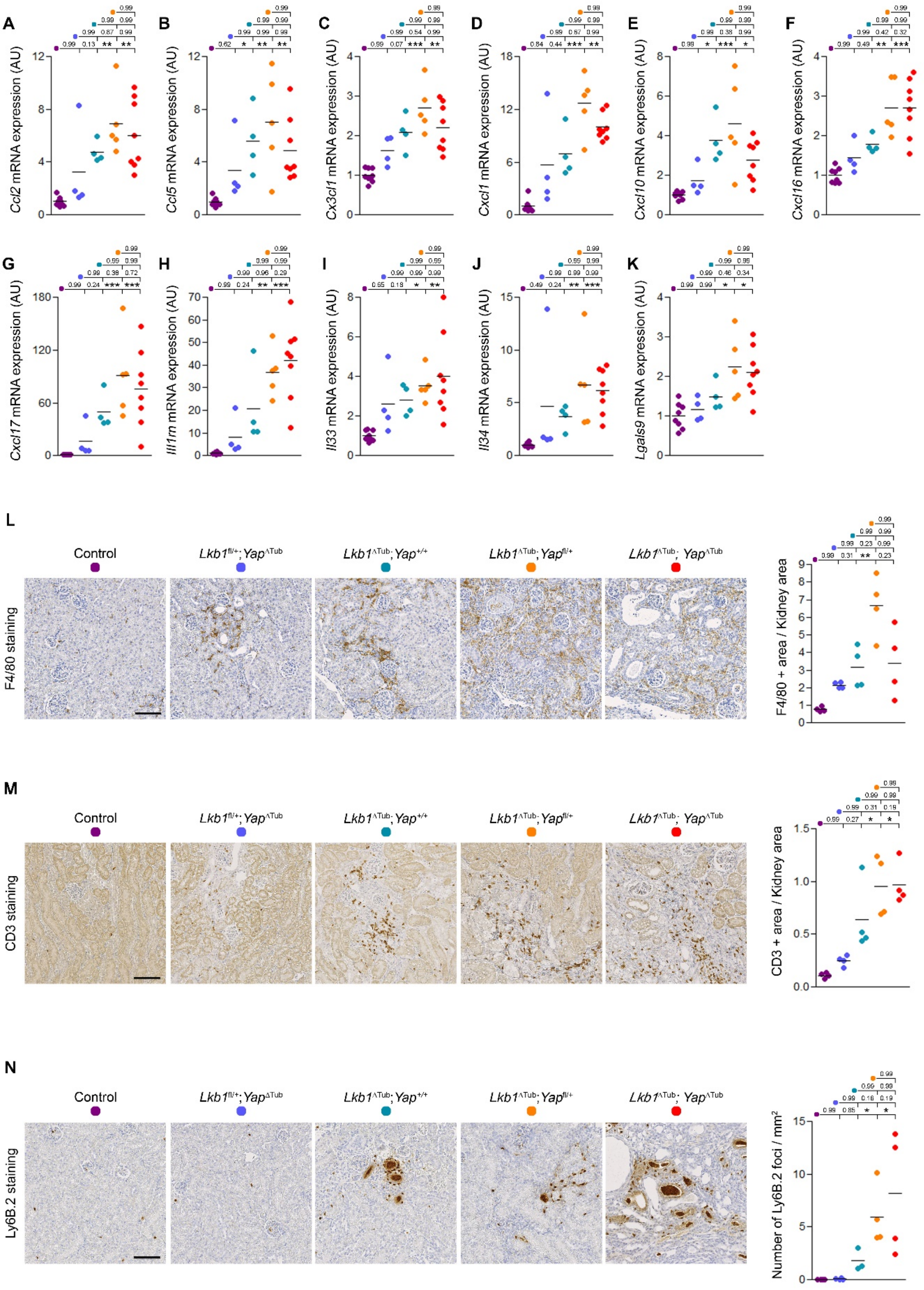


**Supplementary Figure 4. Characterization of renal inflammation after *Yap* depletion in *Lkb1* mutant mice at 8 weeks.**

**(A-K)** Quantification of *Ccl2* (A), *Ccl5* (B), *Cx3cl1* (C), *Cxcl1* (D), *Cxcl10* (E), *Cxcl16* (F), *Cxcl17* (G), *Il1rn* (H), *Il33* (I), *Il34* (J) and *Lgals9* (K) mRNA abundance in kidneys from control, *Lkb1*$^{+/+}$; *Yap*$^{\Delta Tub}$, *Lkb1*$^{fl/+}$; *Yap*$^{\Delta Tub}$, *Lkb1*$^{\Delta Tub}$; *Yap*$^{+/+}$, *Lkb1*$^{\Delta Tub}$; *Yap*$^{fl/+}$ and *Lkb1*$^{\Delta Tub}$; *Yap*$^{\Delta Tub}$ mice at 8 weeks. **(L)** Representative images and quantification of F4/80 (macrophages) immunostaining of kidney sections from 8 weeks old animals. Scale bar: 100µm. **(M)** Representative images and quantification of CD3 (T cells) immunostaining of kidney sections from 8 weeks old mice. Scale bar: 100µm. **(N)** Representative images and quantification of Ly-6B.2 (neutrophils) immunostaining of kidney sections from 8 weeks old mice. Scale bar: 100µm. **(A-N)** Each dot represents one individual mouse. Bars indicate mean. Kruskal-Wallis test, * $P < 0.05$, ** $P < 0.01$, *** $P < 0.001$. AU: arbitrary unit.

**Supplementary Figure 5**

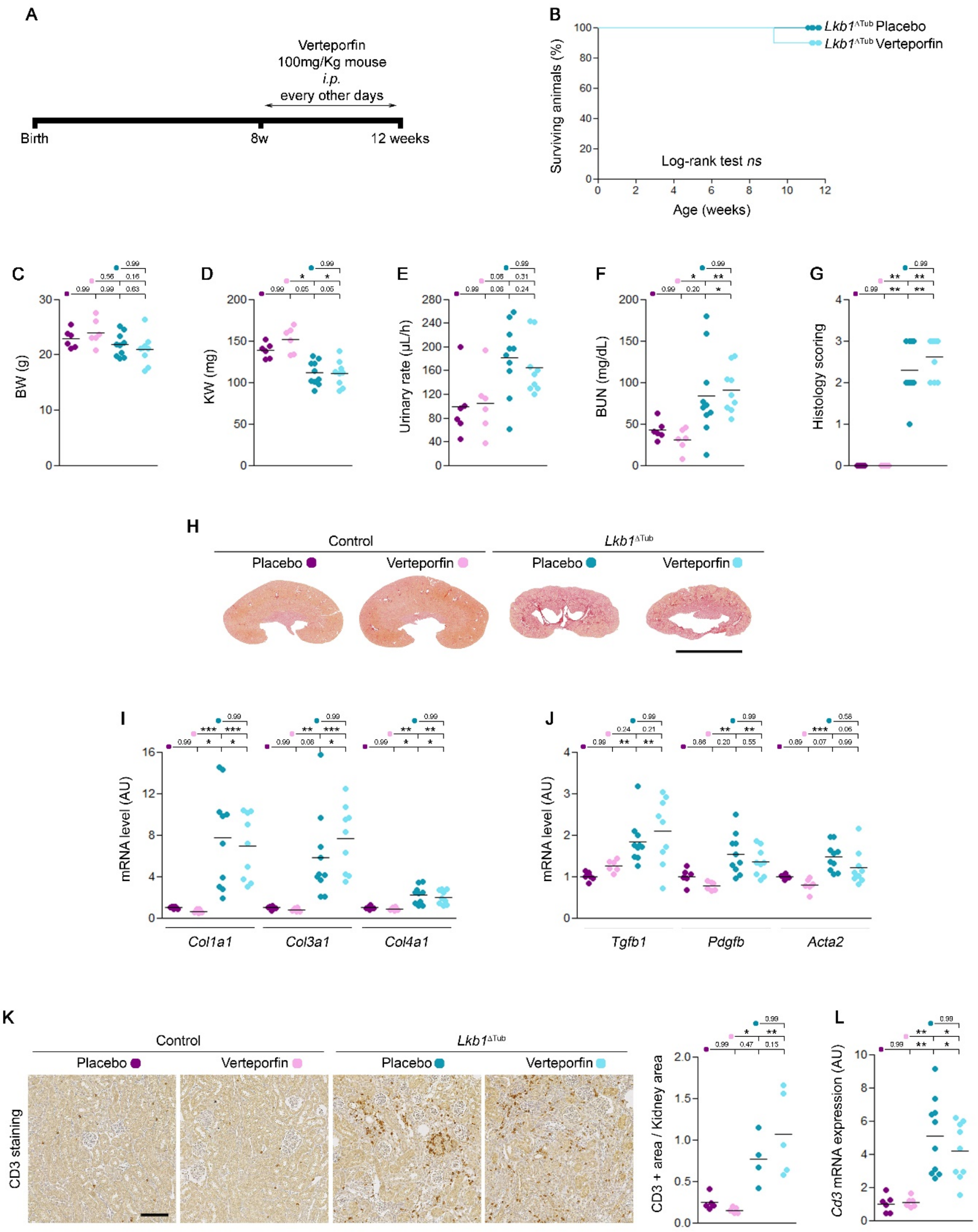


**Supplementary Figure 5. Characterization of pharmacological YAP inhibition in *Lkb1*$^{\Delta Tub}$ mice.**

**(A)** Scheme of verteporfin treatment. Control and *Lkb1*$^{\Delta Tub}$ mice were administered *i.p.* with vehicle (DMSO) or verteporfin (100mg/Kg) every other days from 8 weeks to 12 weeks of age. **(B)** Kaplan–Meier survival curves of the same groups of animals. **(C-F)** Body weight (BW, C), kidney weight (KW, D), urinary rate (E), plasma blood urea nitrogen (BUN, F) at 12 weeks. **(G)** Renal histology lesion scoring at 12 weeks. **(H)** Representative sirius red stained kidney sections at 12 weeks. Scale bars: 5mm. **(I)** Renal collagen mRNA content evaluated by quantitative PCR at 12 weeks. **(J)** *Tgfb1*, *Pdgfb* and *Acta2* mRNA expression in kidneys from the same mice at 12 weeks. **(K)** Representative images and quantification of CD3 (T cells) immunostaining of kidney sections from 8 weeks old mice. Scale bar: 100µm. **(L)** *Cd3* mRNA expression at 12 weeks. **(C-L)** Each dot represents one individual mouse. Bars indicate mean. Kruskal-Wallis test, * $P < 0.05$, ** $P < 0.01$, *** $P < 0.001$. AU: arbitrary unit.

**Supplementary Table 1: Primer pairs used for qRT-PCR.**

| Gene name | Specie | Forward Primer (5' to 3') | Reverse Primer (5' to 3') |
|---|---|---|---|
| *Acta2* | Mouse | CATGCGTCTGGACTTGGCTG | GACAATCTCACGCTCGGCAGTAG |
| *Adgre1* | Mouse | CGTCAGGTACGGGATGAATATAAG | ATCTTGGAAGTGGATGGCATAG |
| *Ankrd1* | Mouse | AGGCTGAACCGCTATAAGATG | GGCTGTCGAATATTGCTTTGG |
| *Areg* | Mouse | TGGTCTTAGGCTCAGGCCATT | TGGTCCCCAGAAAGCGATT |
| *Ccl2* | Mouse | AGTAGGCTGGAGAGCTACAA | GTATGTCTGGACCCATTCCTTC |
| *Ccl5* | Mouse | CCAATCTTGCAGTCGTGTTTG | ACCCTCTATCCTAGCTCATCTC |
| *Cd3* | Mouse | CTGTTCCCAACCCAGACTATG | AAGGCGATGTCTCTCCTATCT |
| *Col1a1* | Mouse | GCCGCAAAGAGTCTACATGTCTAG | TGGCAGATACAGATCAAGCATACC |
| *Col3a1* | Mouse | GGACCAGCAGGAACTAATGGTAT | GTTCTCCAGGTGATCCATCTTT |
| *Col4a1* | Mouse | GTCTGGCTTCTGCTGCTCTTC | CCTTCACGCCATGACAGTCA |
| *Ctgf* | Mouse | GCTGACCTGGAGGAAAACATTAA | TGACAGGCTTGGCGATTTTAG |
| *Cx3cl1* | Mouse | GCTTTGCTCATCCGCTATCA | GTCTTGGACCCATTTCTCCTTC |
| *Cxcl1* | Mouse | CGAAGTCATAGCCACACTCAA | GAGCAGTCTGTCTTCTTTCTCC |
| *Cxcl10* | Mouse | GGCCATAGGGAAGCTTGAAA | CAGACATCTCTGCTCATCATTCT |
| *Cxcl16* | Mouse | ATCAGGTTCCAGTTGCAGTC | CATGACCAGTTCCACACTCTT |
| *Cxcl17* | Mouse | CCTTCCTTCTGTTGCTTCCA | TTCCAAGAGCCACCTCCTA |
| *Cyr61* | Mouse | GGGTTGGAATGCAATTTCGG | TGGTGTTTACAGTTGGGCTG |
| *Edn1* | Mouse | CTGGAGACCCCGCAGGTCCAA | AGTCCATACGGTACGACGCGC |
| *Hprt* | Mouse | GTTAAGCAGTACAGCCCCAAA | AGGGCATATCCAACAACAAACTT |
| *Il1rn* | Mouse | TTGTGCCAAGTCTGGAGATG | CTCAGAGCGGATGAAGGTAAAG |
| *Il33* | Mouse | TGCCTCCCTGAGTACATACA | CTGGTCTTGCTCTTGGTCTTT |
| *Il34* | Mouse | GATATGGACTCTGACCCAAGATAAG | AGCAATCCTGTAGTTGATGGG |
| *Kim1* | Mouse | GAGAGTGACAGTGGTCTGTATTG | CCTTGTAGTTGTGGGTCTTCTT |
| *Lcn2* | Mouse | GGACCAGGGCTGTCGCTACT | GGTGGCCACTTGCACATTGT |
| *Lgals9* | Mouse | CAGATGCTACGAGGTTCCATATC | GTGTTTCGGACAACAGCATTC |
| *Lkb1* | Mouse | CCTGCAAGCAGCAGTGAC | CCAACGTCCCGAAGTGAG |
| *Pdgfb* | Mouse | GGCAAGCACCGAAAGTTTAAG | TAAATAACCCTGCCCACACTC |
| *Ppia* | Mouse | GGCTATAAGGGTTCCTCCTTTC | TTTCTCTCCGTAGATGGACCT |
| *Rpl13* | Mouse | GCTCCAAGCTCATCCTGTT | GGTGGCCAGCTTAAGTTCT |
| *Taz* | Mouse | ACCAAGTACATGAACCACCTC | GTGGTTGGAGACGGTGATAAG |
| *Tgfb1* | Mouse | GGGAAGCAGTGCCCGAACCC | TGGGGGTCAGCAGCCGGTTA |
| *Yap* | Mouse | CCTTTGAGATCCCTGATGATGT | GTTGTTGTCTGATCGTTGTGATT |

| Gene name | Specie | Forward Primer (5' to 3') | Reverse Primer (5' to 3') |
|---|---|---|---|
| *Ankrd1* | Dog | ATCAGTGCTCGGGATAAGTTG | GGTATCTCCTTCTCGGTCTTTG |
| *Hprt* | Dog | AATGTCTTGATTGTTGAGGATAT | ACAAAGTCAGGTTTATAGCC |

**Supplementary Table 2: Antibodies list**
WB: western blot, IHC: immunohistochemistry

#### a. Primary antibodies

| Antibody Name | Company | Species | Application | Dilution | Reference |
|---|---|---|---|---|---|
| $\alpha$-Tubulin | SIGMA | Mouse | WB | 1:10,000 | T5168 |
| FLAG | SIGMA | Mouse | WB | 1:5,000 | F-3165 |
| Lamin A | Abcam | Mouse | WB | 1:100 | ab8980 |
| LATS1 | Cell Signaling Technology | Rabbit | WB | 1:1,000 | 3477 |
| MST1 | Cell Signaling Technology | Rabbit | WB | 1:500 | 3682 |
| YAP/TAZ | Cell Signaling Technology | Rabbit | WB | 1:1,000 | 8418 |
| CD3 | Abcam | Rabbit | IHC | 1:100 | ab16669 |
| F4/80 | Bio-Rad | Rat | IHC | 1:100 | MCA497R |
| LKB1 | Cell Signaling Technology | Rabbit | IHC | 1:100 | 13031 |
| Ly6B.2 | Abcam | Rat | IHC | 1:100 | ab53457 |
| YAP | Cell Signaling Technology | Rabbit | IHC | 1:400 | 14074 |

#### b. Secondary antibodies and other dyes

| Antibody Name | Company | Application | Dilution | Cat. Number |
|---|---|---|---|---|
| Goat anti-Rabbit | Cell Signaling Technology | WB | 1:5,000 | 7074 |
| Goat anti-Mouse | Dako | WB | 1:10,000 | P0447 |
| Rabbit anti-Rat Biotinylated | Vector Laboratories | IHC | 1:200 | BA-4001 |
| Goat anti-Rabbit Biotinylated | Vector Laboratories | IHC | 1:500 | BA-1000 |
| Streptavidin-HRP | Southern Biotech | IHC | 1:2,000 | 7100-05 |
